# Neuroprotective Effects of the Purine-Derived Compound ITH15004 Against Nigericin- and Mitochondria-Induced Toxicity in NSC-34 Motor Neuron-like Cells

**DOI:** 10.64898/2026.11.13.766254

**Authors:** Marta Olazabal, Ricardo de Pacual, Paloma Kênia de Moraes Berenguel Lossavaro, Savannah Ciach, María F. Cano-Abad

## Abstract

Amyotrophic lateral sclerosis (ALS) is a fatal neurodegenerative disorder characterized by the progressive degeneration of motor neurons. Most patients die within 3–5 years of diagnosis, primarily due to respiratory failure, and effective therapeutic options remain limited. Mitochondrial dysfunction, oxidative stress, calcium dysregulation, and neuroinflammation are key contributors to ALS pathogenesis. In this study, we evaluated the potential neuroprotective effects of ITH15004, a novel compound with modulatory effects on P2X7 purinergic receptors, in an in vitro model using NSC-34 motor neuron-like cells. Cellular stress was induced using oligomycin (10 μM) and rotenone (30 μM), inhibitors of mitochondrial oxidative phosphorylation; nigericin (5 μM), a K⁺/H⁺ ionophore that disrupts ionic homeostasis; and extracellular ATP (1 mM), a purinergic receptor agonist. MTT-reducing activity was assessed as an indicator of cellular metabolic activity, while caspase-3/7 activity was measured to evaluate apoptotic responses. Intracellular ATP levels, cytosolic calcium responses, and cytochrome c and VDAC1 protein abundance were also analysed. ITH15004 significantly improved MTT-reducing activity following exposure to ATP, oligomycin/rotenone, and nigericin. In addition, ITH15004 partially restored intracellular ATP levels following nigericin treatment and significantly reduced nigericin-induced caspase-3/7 activation. Western blot analysis further revealed significant changes in cytochrome c and VDAC1 protein levels in response to ITH15004 treatment. These findings indicate that ITH15004 attenuates cellular damage induced by different stressors in NSC-34 cells and modulates several parameters associated with cellular energy status and apoptotic signalling. Further studies are needed to establish its mechanisms of action and assess its therapeutic potential in ALS.

**Highlights:** Nigericin decreases intracellular ATP and increases caspase-3/7 activity in NSC-34 cells.

ITH15004 significantly improves MTT-reducing activity following ATP, oligomycin/rotenone and nigericin exposure.

ITH15004 partially restores intracellular ATP levels after nigericin treatment. ITH15004 significantly reduces nigericin-induced caspase-3/7 activation.

ITH15004 significantly modulates cytochrome c and VDAC1 protein levels in NSC-34 cells.

## Introduction

Amyotrophic lateral sclerosis (ALS) is a progressive neurodegenerative disease characterized by the degeneration of upper motor neurons in the motor cortex and brainstem, and lower motor neurons in the spinal cord. This neuronal loss leads to progressive muscle weakness affecting voluntary movements, speech, swallowing, and ultimately respiratory function, with death typically occurring within 3–5 years of diagnosis due to respiratory failure [1, 2].

ALS exists in two major forms: familial (fALS) and sporadic (sALS). While fALS accounts for approximately 5–10% of cases and follows an autosomal dominant inheritance pattern, sALS represents the majority (90–95%) and lacks a clear genetic cause [3]. However, several genes have been identified as common contributors to both forms of the disease, including *SOD1* (superoxide dismutase 1), *C9orf72* (chromosome 9 open reading frame 72), *TARDBP* (encoding TDP-43), and *FUS* (fused in sarcoma) [3].

ALS pathogenesis involves multiple, converging cellular mechanisms. Among these, glutamate-induced excitotoxicity, oxidative stress, mitochondrial dysfunction, protein aggregation, and neuroinflammatory responses have been strongly implicated [5, 6]. Neuroinflammation is characterized by activation of microglia, reactive astrogliosis, and infiltration of immune cells into the central nervous system. Mitochondrial abnormalities, both structural and functional, are considered central in ALS, contributing to bioenergetic failure, oxidative stress, and the activation of cell death pathways [13]. For instance, mutant *SOD1* proteins accumulate on mitochondrial membranes, disrupting permeability, inhibiting respiratory function, increasing reactive oxygen species (ROS), and inducing the release of cytochrome c—events that promote intrinsic apoptotic signalling [14, 19]. Changes in Bcl-2 family proteins further facilitate mitochondrial membrane permeabilization and the activation of caspases such as caspase-1 and -3, linking both intrinsic and extrinsic apoptotic pathways [19, 22].

Currently approved therapeutic agents—riluzole, edaravone, and relyvrio—offer modest clinical benefits by targeting excitotoxicity, oxidative stress, and mitochondrial/endoplasmic reticulum dysfunction, respectively [2]. The recent approval of tofersen, an antisense oligonucleotide therapy targeting *SOD1* mRNA, provides a personalized option for patients with *SOD1* mutations [23]. Nonetheless, treatment options remain limited, and the identification of novel therapeutic strategies remains a critical unmet need.

Spinal motor neurons display selective vulnerability in ALS, attributed in part to their high expression of Ca²⁺-permeable AMPA receptors and poor calcium-buffering capacity, making them especially sensitive to ionic dysregulation and mitochondrial stress [Van Den Bosch et al., 2006]. Consequently, the elucidation of mitochondrial contributions to apoptotic signalling pathways is essential for a comprehensive understanding of ALS pathophysiology.

Amyotrophic lateral sclerosis (ALS) is a progressive neurodegenerative disease characterized by the degeneration of upper motor neurons in the motor cortex and brainstem, and lower motor neurons in the spinal cord. This neuronal loss leads to progressive muscle weakness affecting voluntary movements, speech, swallowing, and ultimately respiratory function, with death typically occurring within 3–5 years of diagnosis due to respiratory failure [1, 2].

ALS exists in two major forms: familial (fALS) and sporadic (sALS). While fALS accounts for approximately 5–10% of cases and follows an autosomal dominant inheritance pattern, sALS represents the majority (90–95%) and lacks a clear genetic cause [3, 4]. However, several genes have been identified as common contributors to both forms of the disease, including SOD1 (superoxide dismutase 1), C9orf72 (chromosome 9 open reading frame 72), TARDBP (encoding TDP-43), and FUS (fused in sarcoma) [3].

ALS pathogenesis involves multiple, converging cellular mechanisms. Among these, glutamate-induced excitotoxicity, oxidative stress, mitochondrial dysfunction, protein aggregation, and neuroinflammatory responses have been strongly implicated [5–7]. Neuroinflammation is characterized by activation of microglia, reactive astrogliosis, and infiltration of immune cells into the central nervous system [6, 7]. Mitochondrial abnormalities, both structural and functional, are considered central in ALS, contributing to bioenergetic failure, oxidative stress, and the activation of cell death pathways [7–9]. For instance, mutant SOD1 proteins can associate with mitochondrial membranes, disrupting mitochondrial function, increasing reactive oxygen species (ROS) production, and promoting cytochrome c release, thereby contributing to intrinsic apoptotic signalling [8–10]. Mitochondrial damage can also trigger caspase activation and amplify apoptotic pathways, including caspase-1- and caspase-3-mediated cell death [10–12].

Currently approved therapeutic agents, including riluzole and edaravone, offer modest clinical benefits by targeting excitotoxicity and oxidative stress, respectively [2, 13]. Tofersen, an antisense oligonucleotide therapy targeting SOD1 mRNA, provides a personalized therapeutic option for patients carrying SOD1 mutations [14]. Nonetheless, treatment options remain limited, and the identification of novel therapeutic strategies remains a critical unmet need.

Spinal motor neurons display selective vulnerability in ALS, attributed in part to their high expression of Ca²⁺-permeable AMPA receptors and poor calcium-buffering capacity, making them especially sensitive to calcium dysregulation and mitochondrial stress [15]. Consequently, elucidating the contribution of mitochondrial dysfunction to apoptotic signalling pathways is essential for a comprehensive understanding of ALS pathophysiology.

In this study, we employed the NSC-34 motor neuron-like cell line, a widely used in vitro model for investigating motor neuron-related mechanisms and neurodegenerative processes [16, 23], to investigate cytotoxic mechanisms relevant to ALS. We exposed cells to several insults: (i) nigericin, a potassium ionophore that induces K⁺ efflux and disrupts intracellular ionic homeostasis, mimicking certain downstream consequences associated with P2X7 receptor activation; (ii) extracellular ATP, to activate purinergic receptors; and (iii) a combination of oligomycin and rotenone (O/R), to induce mitochondrial stress through inhibition of mitochondrial respiratory complexes [17–20].

We further evaluated the neuroprotective effects of ITH15004, a novel purine derivative synthesized by Calzaferri et al. and characterized as a P2X7 receptor antagonist [21]. In addition to modulating purinergic signalling, ITH15004 regulates mitochondrial Ca²⁺ homeostasis and enhances exocytotic activity in chromaffin cells, suggesting a multifaceted mechanism of action [22]. These properties position ITH15004 as a promising candidate for modulating mitochondrial dysfunction and purinergic signalling pathways in ALS.

### Materials and Methods Cell Culture

NSC-34 cells, a murine motor neuron-like hybrid cell line, were used throughout the study. Cells were maintained in a humidified incubator at 37 °C under a 5% CO atmosphere in a filter-sterilized culture medium consisting of DMEM high glucose (13.5 g/L) supplemented with 10% fetal bovine serum (FBS), 3.7 g/L NaHCO, 1.19 g/L HEPES and 5 mL penicillin/streptomycin (25 IU/mL penicillin, and 25 µg/mL streptomycin). Cells were routinely passaged at sub-confluence and used for functional experiments between passages 14 and 24.

### Immunocytochemistry for P2X7 Receptor Detection

NSC-34 cells seeded on glass coverslips were washed with phosphate-buffered saline (PBS) and fixed with 3.5% paraformaldehyde in PBS for 20 min at room temperature. Cells were permeabilized with 0.1% Triton X-100 for 15 min and blocked with 10% foetal bovine serum (FBS) in PBS for 30 min. Samples were incubated with a rabbit anti-P2X7 primary antibody (1:200, Cat# APR-004, Alomone Labs) for 45 min at 100 rpm, followed by a secondary goat anti-rabbit IgG antibody (1:500, Cat# sc-2004, Santa Cruz Biotechnology) under the same conditions. Nuclei were counterstained with DAPI (1:500), and coverslips were mounted using fluorescence-compatible media. Images were captured using a Leica TCS SP5 confocal microscope equipped with 405 nm, 488 nm, and 561 nm laser lines and a 63× oil immersion objective.

### MTT Cell Viability Assay

MTT-reducing activity was evaluated using the MTT colorimetric assay. Rapid colorimetric assay for cellular growth and survival: application to proliferation and cytotoxicity assays. in NSC-34 cells under the toxic paradigms: oligomycin (10 µM)/rotenone (30 µM), nigericin (5 µM), or ATP (1 mM)[24].

NSC-34 cells were seeded in 48-well plates at 5×10 cells/well. After 24 h, cells were pretreated with ITH15004 for 24 h, then co-treated with cytotoxic agents. The co-incubation period was 24 h for all treatments, except for ATP (48 h). The endpoint was defined a priori as MTT-reducing activity and not as a direct measure of cell number, membrane integrity, cytoprotection, or neuroprotection.

MTT (0.5 mg/mL) was added to the wells and incubated for 15 min at 37 °C. Formazan was solubilized with DMSO and absorbance was measured at 540 nm using a FLUOstar Optima reader (BMG Labtech, Ortenberg, Germany). Results were expressed as percentage of MTT- reducing activity relative to the corresponding untreated control, set to 100%. Replicate wells were performed within each independent culture. For inferential statistics, replicate wells from the same culture and experimental condition were averaged to obtain one value per culture, and the independent culture was used as the biological unit. MTT-reducing activity was calculated as: (OD_test/OD_control) × 100%.

### Caspase-3/7 Activity Assay

Caspase activation was measured using the Caspase-Glo® 3/7 assay kit (Promega). NSC-34 cells were seeded at 5×10 cells/well in 96-well plates. Cells were pretreated with ITH15004 for 24 h, followed by exposure to nigericin (5 µM) for 24 h. Equal volumes of Caspase-Glo reagent (100 µL/well) were added, and luminescence was measured after 30 min incubation at room temperature using a luminometer (Promega). The optimal incubation time was determined from preliminary time-course experiments (Supplementary Figure S2).

### ATP Quantification Using CellTiter-Glo®

Intracellular ATP levels were quantified using the CellTiter-Glo® luminescent assay (Promega). Cells were seeded at 5×10 cells/well in 96-well plates, pretreated with ITH15004 for 24 h, and then incubated with nigericin (5 µM) for an additional 24 h. After treatment, equal volumes of CellTiter-Glo reagent were added, mixed for 2 min on an orbital shaker, and incubated for 10 min at room temperature. Luminescence was recorded using a luminometer. A separate set of experiments was used for baseline cytotoxicity (Supplementary Figure S1).

### Intracellular Ca²⁺ Imaging with Fura-2

Calcium imaging was performed using the ratiometric indicator Fura-2 AM. Cells on coverslips were washed and incubated with 2.5 µM Fura-2 AM in Krebs-HEPES solution (composition in mM: 145 NaCl, 5.9 KCl, 1.2 MgCl, 2 CaCl, 10 HEPES, 10 D-glucose; pH 7.4) for 45 min at 37°C and 5% CO . Coverslips were mounted in a perfusion chamber and imaged using an Olympus BX51WI microscope with a 60× water immersion objective. Excitation was alternated between 340 and 380 nm, with emission collected at 510 nm. Images were captured every 2 s using a CCD camera and analyzed with Olympus Xcellence software.

### Western blot

#### Western blot analysis of cytochrome c and VDAC1

NSC-34 motor neuron-like cells were seeded in 6-well plates (2 × 10 cells/well) and treated for 24 h with vehicle (control), nigericin (5 µM), ITH15004 (3 µM), or the combination of both drugs. After treatments, cells were washed twice with ice-cold phosphate-buffered saline (PBS) and lysed on ice in RIPA buffer (50 mM Tris-HCl, 150 mM NaCl, 1 mM EDTA, 1% NP-40, 0.1% SDS, pH 7.4) supplemented with protease and phosphatase inhibitors (Roche). Lysates were centrifuged at 12,000 × g for 15 min at 4 °C, and the supernatants were collected for protein quantification using the BCA assay (Thermo Fisher).

Equal amounts of protein (20–25 µg) were separated on 10–12% SDS–polyacrylamide gels and transferred onto PVDF membranes (0.45 µm, Millipore) using a semi-dry transfer system. Membranes were blocked for 1 h at room temperature in 5% non-fat dry milk prepared in Tris-buffered saline containing 0.1% Tween-20 (TBST) and subsequently incubated overnight at 4 °C with the following primary antibodies:

- anti–cytochrome c (1:1000; Cell Signaling Technology),
- anti–VDAC1 (1:2000; Abcam, ab15895), and
- anti–GAPDH (1:5000; Sigma-Aldrich) as loading control.

After washing with TBST, membranes were incubated for 1 h at room temperature with HRP-conjugated secondary antibodies (1:5000; Jackson ImmunoResearch). Protein bands were visualized using an enhanced chemiluminescence (ECL) substrate (Thermo Fisher) and imaged with a ChemiDoc MP Imaging System (Bio-Rad).

For densitometric quantification, band intensities were analyzed using ImageJ (NIH). Cytochrome c signals were normalized either to GAPDH (for cytosolic loading control) or to VDAC1 (for mitochondrial mass correction). Data were expressed as relative optical density (arbitrary units) and averaged from three to four independent experiments. Statistical analyses were performed using one-way ANOVA followed by Tukey’s post hoc test, with significance set at *p < 0.05*.

### Reagents

NSC-34 cells were obtained from ATCC. ITH15004 was synthesized at the Teófilo Hernando Institute for Drug Discovery (Madrid, Spain). Nigericin was purchased from MedChemTronica. Other reagents (DMEM, FBS, P/S, trypsin/EDTA, oligomycin, rotenone, ATP) were from Sigma-Aldrich. rabbit anti-P2X7 primary antibody (1:200, Cat# APR-004, Alomone Labs), secondary goat anti-rabbit IgG antibody (1:500, Cat# sc-2004, Santa Cruz Biotechnology) Caspase-Glo® and CellTiter-Glo® kits were from Promega.

### Statistical Analysis

Data are presented as single examples of the original recording to illustrate the experimental protocols and as averaged pooled data from several experiments, as means ± SEM of the number of different experiments done (number of cells, n) and different cultures used (N).

Statistical analyses were done using the GraphPad Prism, version 8.0 (GraphPad Software Inc. La Jolla, CA, USA). Student’s paired t-test was used only for a single prespecified two-condition within-culture comparison. When three or more concentrations or sequential treatment conditions were evaluated within the same cultures, one-way repeated-measures ANOVA was used, followed, when planned contrasts were required, by Tukey multiple comparison test post-hoc. The level of statistical significance was established at p<0.05.

## Results

### Expression of P2X7 Receptor in NSC-34 Cells

ITH15004 was synthesized as a P2X7 receptor antagonist, as described by Calzaferri et al. (2021). Immunofluorescence analysis demonstrated that NSC-34 motor neuron- like cells express P2X7 receptors. Confocal microscopy revealed prominent green fluorescence corresponding to P2X7R (Fig. 1a–b). Cell nuclei were successfully stained with DAPI (blue) and colocalized with cellular structures (Fig. 1a).

**Figure 1.**
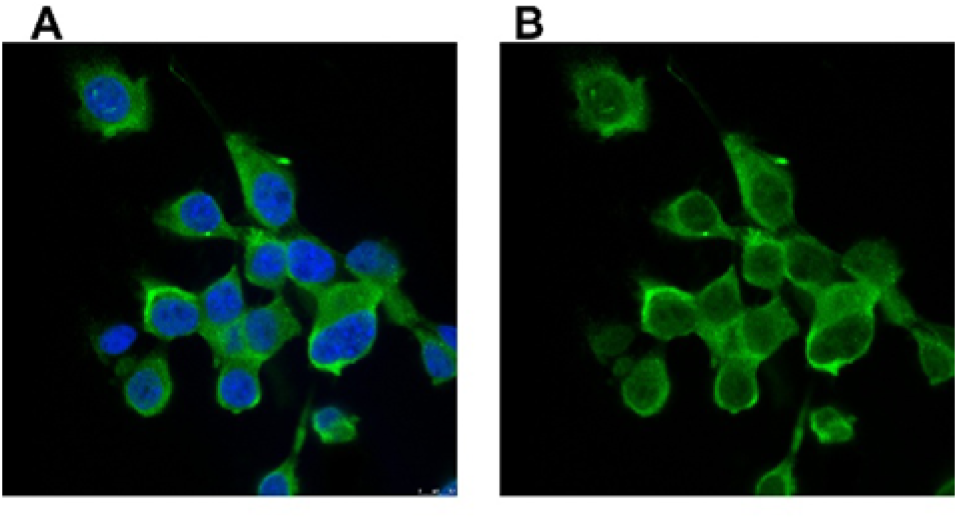
Immunofluorescence of P2X7R (green) and nuclei (blue) in NSC-34 cells. Highly fluorescent green represents receptor expression.

### Immunofluorescence analysis of P2X7 receptor expression in NSC-34 cells

Immunofluorescence analysis using confocal microscopy demonstrated that NSC-34 motor neuron-like cells express P2X7 receptors on their surface. The cells were incubated with a specific primary antibody against P2X7R, followed by a fluorescently labeled secondary antibody. As shown in Figure 1a–b, prominent green fluorescence corresponding to P2X7R immunolabeling was detected, predominantly localized at the plasma membrane.

Cell nuclei were counterstained with DAPI (blue), allowing for clear visualization of nuclear morphology and identification of individual cells. The DAPI signal colocalized with the cell body, and the membrane-localized green fluorescence indicates the surface expression of P2X7 receptors in NSC-34 cells. These findings support the presence of functional P2X7R at the cell membrane, potentially involved in purinergic signaling pathways in this motor neuron-like cell line.

## ITH15004 preserves MTT-reducing activity in NSC-34 Cells exposed to

### Oligomycin/Rotenone, ATP, and Nigericin

To model mitochondrial dysfunction and oxidative stress in ALS, NSC-34 cells were treated with oligomycin/rotenone (O/R, 10/30 µM) for 24 h. Cell viability, assessed via MTT assay, was significantly reduced by 49 ± 9.4% (Fig. 2a). No further cytotoxicity was observed at higher concentrations (data not shown).

**Figure 2.**
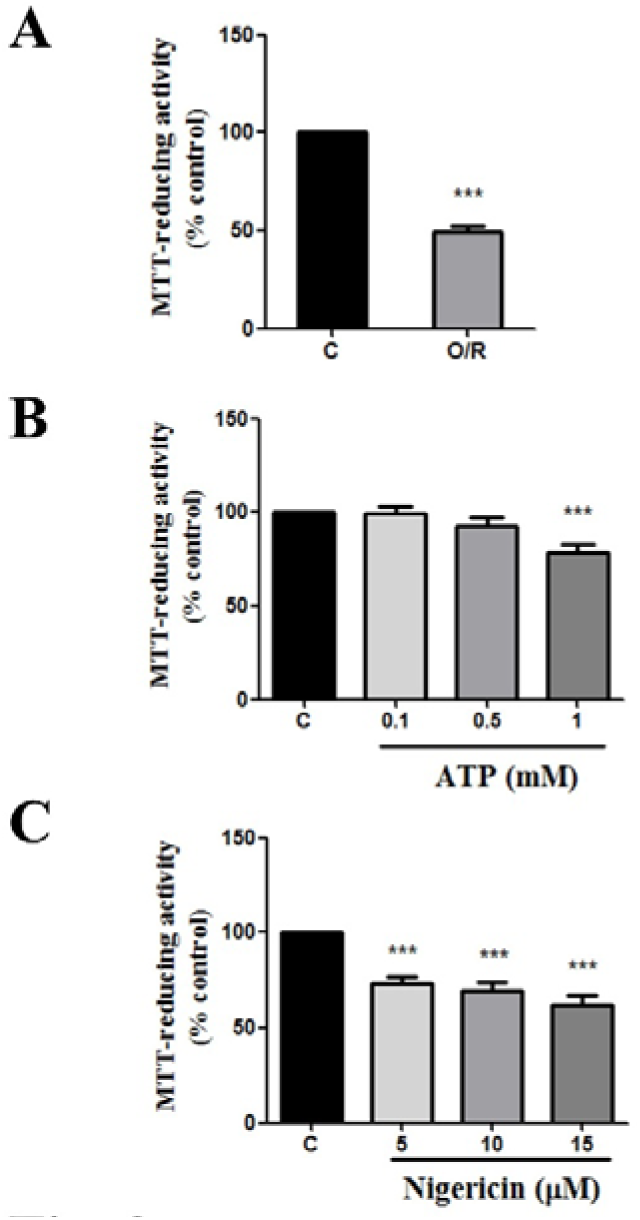
Cell viability of NSC-34 cells following (a) O/R (10/30 µM, 24 h), (b) ATP (0.1–1 mM, 48 h), and (c) Nigericin (5–15 µM, 24 h). The treatment is displayed at the top. Data are expressed as mean ± SEM; N=3–4; n=9–12; Student’s paired t-test was used in panel A (t= 20.81; df= 32; R^2^= 0.9321; ***p<0.0001); one-way repeated- measures ANOVA, followed by Tukey multiple comparison test post-hoc (F=13.28; R^2^= 0.6241; ***p<0.000, for panel B and F=27.09; R^2^=0.6930; ***p<0.0001, for panel C).

ATP induced a dose-dependent reduction in viability after 48 h of exposure (0.1–1 mM), with a significant 21.4 ± 5.8% decrease at 1 mM (Fig. 2b), consistent with P2X7 receptor activation.

Nigericin (5–15 µM, 24 h) induced significant cytotoxicity (viability: 74 ± 2.9%, 73.4 ± 3.3%, and 71.9 ± 4.1%, respectively; Fig. 2c), with no dose-dependent effect. Thus, 5 µM was selected for further assays.

### Neuroprotective Effects of ITH15004 Against Cytotoxic Insults

ITH15004 (1–10 µM) showed no intrinsic cytotoxicity modified the MTT-reducing activity after 24 h (Supplementary Fig. S1). For neuroprotection assays, NSC-34 cells were pretreated with ITH15004 (3 µM, 24 h), followed by co-incubation with O/R, ATP, or Nigericin for an additional 24 h.

ITH15004 significantly improved viability against and restored MTT-reducing activity viability in ATP-treated cells from 70.1 ± 4.6% to 77.8 ± 4.0% (Fig. 3a) and O/R (50.5 ± 9.5%; Fig. 3b), Remarkably, ITH15004 reversed Nigericin-induced toxicity (46.5 ± 5.6%, Fig 3c), increasing viability restored MTT-reducing activity to levels control (103 ± 4.0%) (Fig. 3c).

**Figure 3.**
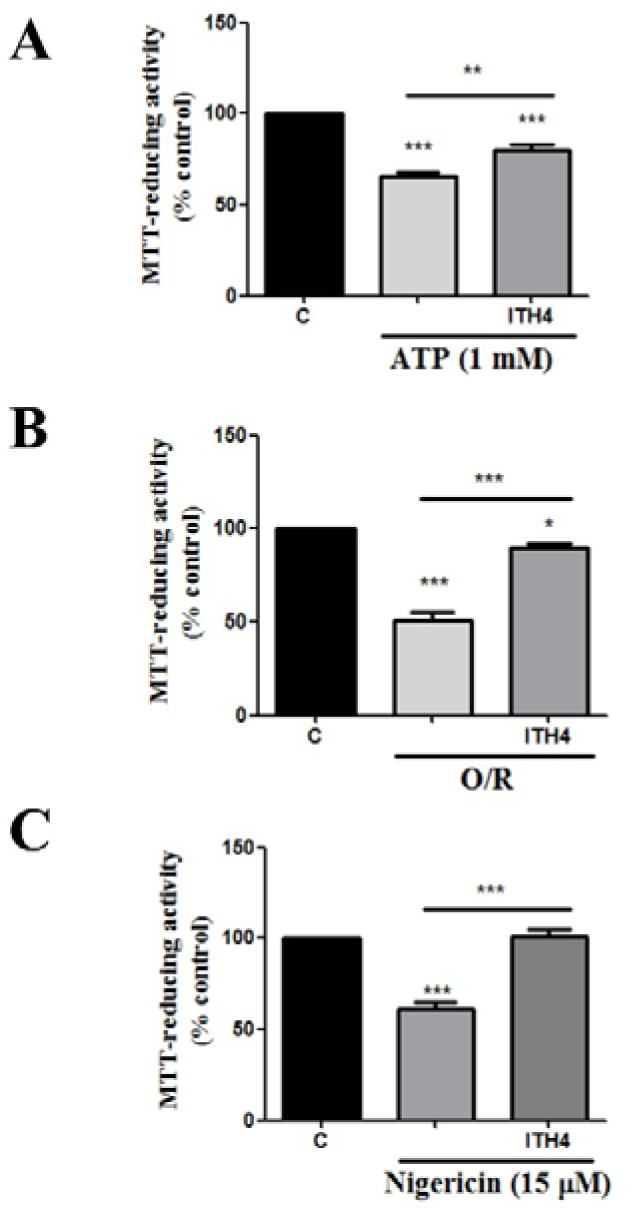
Cell viability of NSC-34 cells pretreated with ITH15004 (3 µM, 24 h) followed by co-treatment with (a) O/R, (b) ATP (1 mM), and (c) Nigericin (5 µM, 24 h). The treatment is displayed at the top. N=4; n=12; one-way repeated-measures ANOVA, followed by Tukey multiple comparison test post-hoc, (F=118.0; R^2^=0.9147; ***p<0.0001respect to control and **p<0.005 respect to ATP vs ATP+ITH15004 for panel A; F=92.46; R^2^= 0.8937; ***p<0.0001 control vs O/R treatment; *p<0.05 control vs O/R + ITH15004 and ***p<0.0001 O/R vs O/R + ITH15004 for panel B and F= 75.69; R^2^= 0.9044; ***p<0.0001 Nigericin vs control and ***p>0.0001 Nigericin vs Nigericin + ITH15004 for panel C),

### Caspase-3/7 Activation and ITH15004 Neuroprotection

To assess apoptosis, caspase-3/7 activation was measured after treatment with O/R, ATP, or Nigericin (5 µM, 24 h). Peak activation occurred at 0.5 h post-treatment (Supplementary Fig. S2). O/R augmented caspase-3/7 activation significantly (163.39 ± 14.3%); ATP reduced the activation of caspase-3/7 to 51.5 ± 1.8% and Nigericin induced a significantly higher caspase-3/7 activation (356 ± 0.2%; 310.3 ± 23.3 % Fig. 4a).

**Figure 4.**
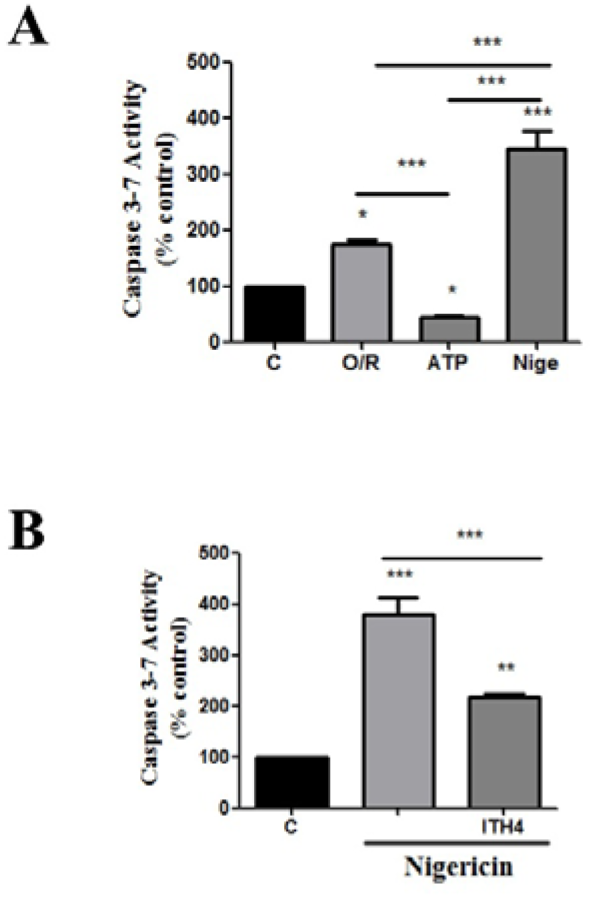
(a) Caspase-3/7 activation in NSC-34 cells following 24 h exposure to cytotoxic agents. (b) ITH15004 (3 µM) pretreatment attenuates Nigericin-induced caspase activation. N=3; n=9. The treatment is displayed at the top. One-way repeated-measures ANOVA, followed by Tukey multiple comparison test post-hoc, (F= 52.11; R^2^= 0.9287; ***p<0.0001 control vs Nigericin; O/R vs ATP; O/R vs Nigericin and ATP vs Nigericin and *p< 0.05 control vs O/R and C vs ATP for panel A and F=55.68; R^2^= 0.9176; ***p<0.0001 control vs Nigericin and Nigericin vs Nigericin+ITH15004 and **p<0.005 control vs Nigericin+ITH15004 for panel B).

ITH15004 pretreatment (3 µM, 24 h) followed by co-exposure to Nigericin reduced caspase activity from 343 ± 28.6% 310.3 ± 23.3% to 258 ± 56% 217.71 ± 17.4%, supporting its anti-apoptotic properties (Fig. 4b).

### ATP Production and Mitochondrial Preservation by ITH15004

O/R and ATP did not show a significant reduction in remaining ATP. Nigericin significantly reduced ATP levels to 36.8 ± 2.2% (Fig. 5a). ITH15004 pretreatment partially restored ATP production to 65.9 ± 4.3% (Fig. 5b), indicating mitochondrial preservation.

**Figure 5.**
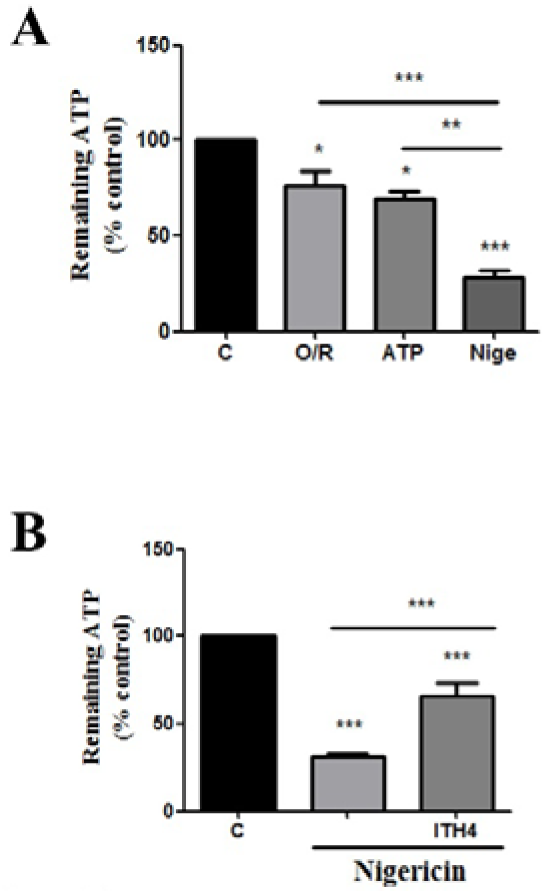
ATP quantification following (a) cytotoxic treatments and (b) ITH15004 pretreatment + Nigericin. N=3; n=9; Student’s t-test, p < 0.05. The treatment is displayed at the top. One-way repeated-measures ANOVA, followed by Tukey multiple comparison test post-hoc, (F= 32.04; R^2^= 0.9144; ***<p=0.0001 control vs Nigericin and O/R vs Nigericin; *p<0.05 control vs O/R and control vs ATP and **p<0.005 ATP vs Nigericin for panel A; F=60.95; R^2^= 0.8840; ***p<0.0001 control vs Nigericin; control vs Nigericin + ITH15004 and Nigericin vs Nigericin + ITH15004 for panel B)

### Cytosolic Ca²⁺ Dysregulation and ITH15004 Modulation

Nigericin induced a concentration-dependent elevation in cytosolic Ca²⁺ ([Ca²⁺]c), measured using Fura-2, with the highest increase observed at 15 µM (Fig. 6A–B). Representative Fura-2 fluorescence recording showing the increase in cytosolic Ca²⁺ ([Ca²⁺]c) following sequential administration of nigericin at 5, 10, and 15 µM, after a 2- min baseline recording (figure 6A). Figure 6B, shows the bar graph the quantification of cytosolic Ca²⁺ amplitude (AFU) at the different nigericin concentrations compared with basal levels. Nigericin induced a concentration-dependent increase in cytosolic Ca²⁺, reaching its highest amplitude at 15 µM.

**Figure 6.**
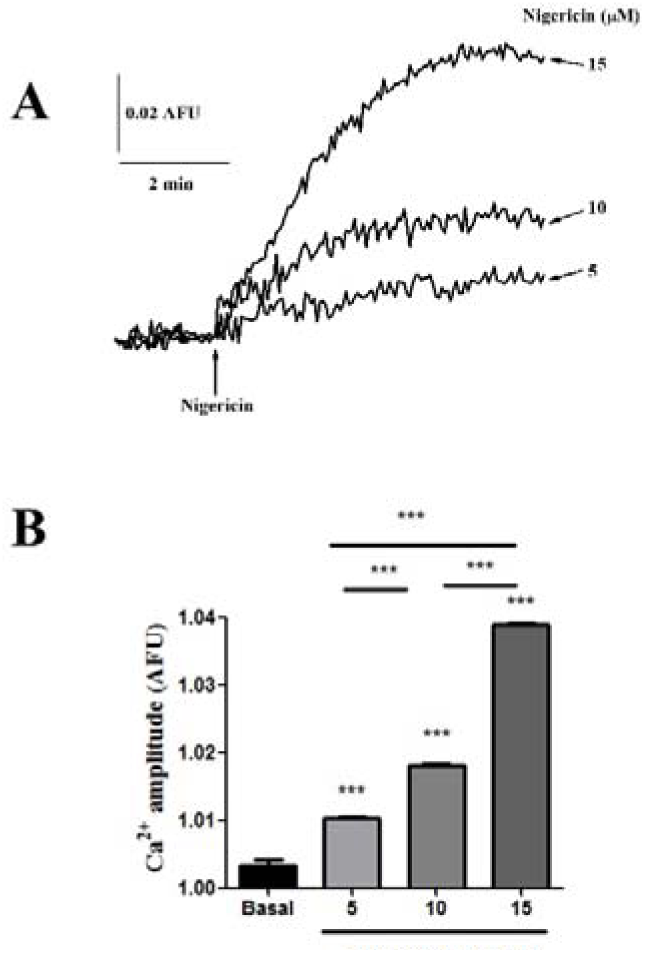
(a) Representative original [Ca²⁺]c traces in NSC-34 cells perfused with Nigericin (5–15 µM, 8 min). (b) AUC AFU quantification of [Ca²⁺]c changes respect to basal line (before nigericin introduction). N=5–13; n=18–82. The treatment is displayed at the top. One-way repeated-measures ANOVA, followed by Tukey multiple comparison test post-hoc, (F= 1282; R^2^= 0.9930; ***<p=0.0001 for all condition in panel B)

### ITH15004 Attenuates Nigericin-Induced Cytosolic Ca²⁺ Elevation in NSC-34 Cells

Nigericin induced an increase in cytosolic Ca²⁺ ([Ca²⁺]c) in NSC-34 cells, which was attenuated by co-incubation with ITH15004 (ITH15004). Notably, the Ca²⁺ response elicited by nigericin was lower than the global Ca²⁺ response induced by subsequent ATP application, indicating a differential magnitude of intracellular Ca²⁺ mobilization. These findings suggest that ITH15004 modulates the nigericin-induced increase in cytosolic Ca²⁺, attenuating Ca²⁺ entry or accumulation in the cytosol.

Experiments evaluating NSC-34 proliferation with ITH15004 and treatments with LPS did not yield significant changes in viability (data not shown).

### ITH15004 preserves mitochondrial integrity and increases mitochondrial mass in NSC-34 cells

To assess whether ITH15004 protects mitochondrial integrity under stress conditions, cytochrome C and VDAC1 protein levels were analysed by Western blot in NSC-34 cells (Figure 8A). Treatment with nigericin (5 µM) markedly decreased cytochrome c levels compared with control cells, consistent with mitochondrial outer-membrane permeabilization and release of pro-apoptotic factors. In contrast, exposure to ITH15004 (3 µM) alone increased cytochrome c abundance, suggesting enhanced mitochondrial content or biogenesis. Notably, co-treatment with ITH15004 and nigericin restored cytochrome c expression to near-control levels.

**Figure 7.**
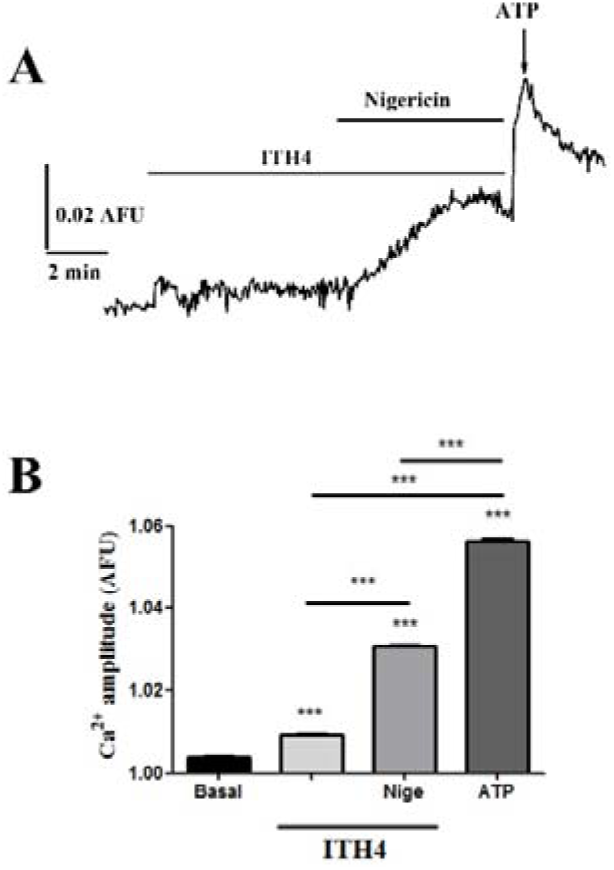
(a) Representative original Ca²⁺ trace with ITH15004 (3 µM), followed by ITH15004 + Nigericin (15 µM), then ATP (1 mM). (b) AUC comparison. N=6; n=39. The treatment is displayed at the top. One-way repeated-measures ANOVA, followed by Tukey multiple comparison test post-hoc, (F= 2870; R^2^= 0.9969; ***<p=0.0001 for all condition in panel B)

**Figure 8.**
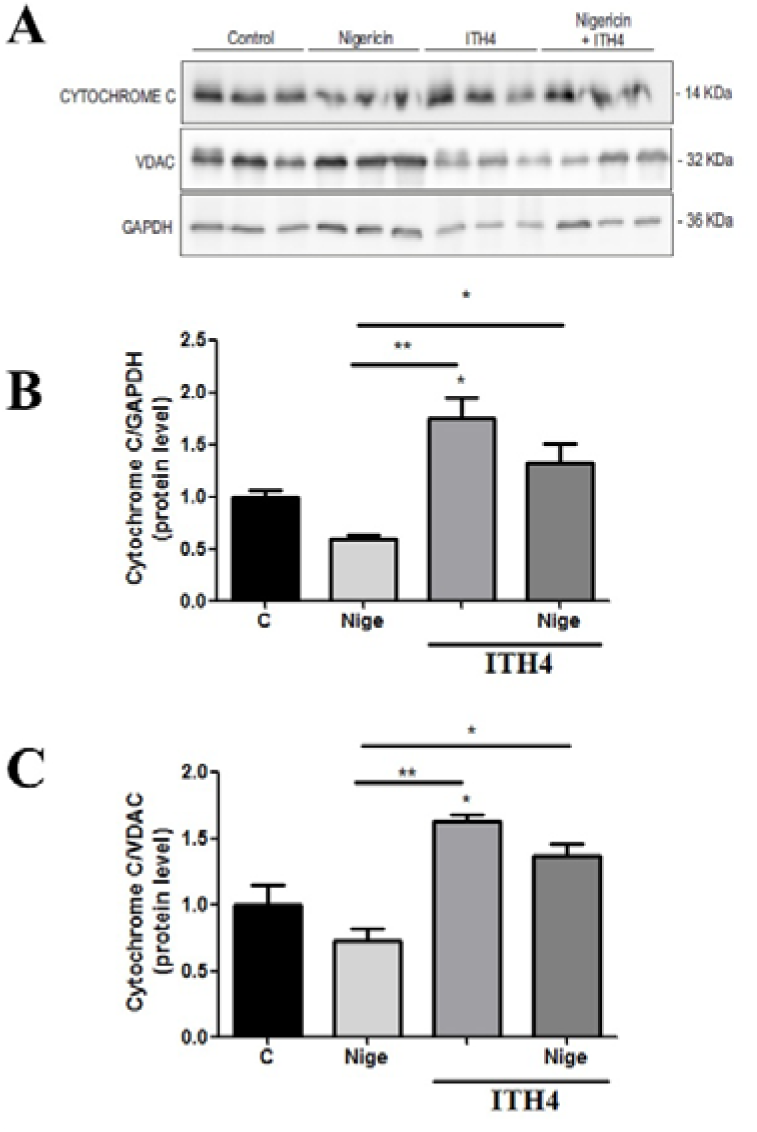
ITH15004 preserves mitochondrial integrity and cytochrome c content in NSC-34 cells. (A) Representative Western blots showing cytochrome c, VDAC1, and GAPDH levels after treatment with vehicle (Control), nigericin (5 µM), ITH15004 (3 µM), or their combination. Data are presented as mean ± SEM (n = 3–4 independent experiments). The treatment is displayed at the top. B) Densitometric analysis of cytochrome c normalized to GAPDH. One-way repeated-measures ANOVA, followed by Tukey multiple comparison test post-hoc, (F= 12.33; R^2^= 0.8604; **p=0.00056 for Nigericin vs ITH15004+Nigericin; *P<0.05 control vs ITH15004 and nigericin vs nigericin +ITH15004 for panel B). (C) Ratio of cytochrome c to VDAC1. One-way repeated-measures ANOVA, followed by Tukey multiple comparison test post-hoc, (F= 12.13; R^2^= 0.8585; **p=0.00059 for Nigericin vs ITH15004+Nigericin; *P<0.05 control vs ITH15004 and nigericin vs nigericin +ITH15004 for panel C).

Densitometric quantification normalized to GAPDH confirmed the significant reduction of cytochrome c induced by nigericin and its reversal by ITH15004 co-incubation (Figure 8B). To better estimate mitochondrial mass, VDAC1—a mitochondrial outer- membrane protein and established mitochondrial loading control—was also evaluated. The pattern of VDAC1 expression paralleled that of cytochrome c, showing decreased signal in nigericin-treated cells and increased levels after ITH15004 exposure. When the ratio of cytochrome c to VDAC1 was calculated (Figure 8C), nigericin significantly lowered this index, whereas ITH15004 alone elevated it and co-treatment normalized it to control values. These results indicate that ITH15004 counteracts the nigericin- induced loss of mitochondrial proteins and supports mitochondrial biogenesis or stability in NSC-34 cells.

## Discussion

Mitochondrial dysfunction, altered calcium homeostasis and neuroinflammation are recognised contributors to amyotrophic lateral sclerosis (ALS) pathophysiology [6–9,15,27]. These processes may converge to compromise motor neuron function and survival, highlighting the importance of identifying compounds capable of limiting cellular damage under conditions of mitochondrial and ionic stress [2,5]. In the present study, we investigated the effects of the purine-derived compound ITH15004 (ITH15004) in NSC-34 motor neuron-like cells exposed to different cytotoxic stimuli, including oligomycin/rotenone (O/R), extracellular ATP and nigericin. Our results show that ITH15004 improves MTT-reducing activity under these conditions and attenuates several responses associated with nigericin-induced cellular damage, including ATP depletion and caspase-3/7 activation. In addition, changes in cytosolic calcium and mitochondrial protein abundance provide further information about the cellular responses to nigericin and ITH15004.

NSC-34 cells are a widely used motor neuron-like cell model for investigating mechanisms associated with neuronal injury and neurodegeneration [16,23]. Although this cell line cannot reproduce the complexity of ALS pathology in vivo, it provides a useful experimental system for studying cellular responses to defined stressors and evaluating potential protective compounds.

In our experiments, O/R, ATP and nigericin produced distinct effects on MTT-reducing activity. O/R, a combination of oligomycin and rotenone, caused a marked reduction in this parameter. This response is consistent with the established pharmacological actions of these compounds: rotenone inhibits mitochondrial complex I, whereas oligomycin blocks ATP synthase, thereby interfering with mitochondrial oxidative phosphorylation [19,20]. These agents provide an experimental approach to investigate cellular responses to impaired mitochondrial energy metabolism, a process implicated in ALS [7–9].

Extracellular ATP also reduced MTT-reducing activity in a concentration-dependent manner, with a significant effect at 1 mM. ATP is an extracellular signalling molecule that activates purinergic receptors, including P2X7, which have been implicated in neuroinflammation and neuronal injury [17,18]. Previous studies have shown that P2X7 receptor activation can contribute to neuronal cell death through caspase-dependent mechanisms [12]. Importantly, our immunofluorescence experiments detected P2X7 receptor immunoreactivity in NSC-34 cells, providing evidence of receptor expression in this cellular model. However, receptor expression alone does not establish its functional contribution to the responses observed following ATP exposure.

Nigericin also significantly reduced MTT-reducing activity in NSC-34 cells. This compound is a K⁺/H⁺ ionophore that alters transmembrane ion gradients [28]. It is widely used experimentally to promote potassium efflux and activate the NLRP3 inflammasome in immune cells, through a mechanism that does not require direct activation of P2X7 receptors [25]. Therefore, although extracellular ATP and nigericin can both produce ionic disturbances, their mechanisms of action are distinct. This distinction is relevant when interpreting the effects of ITH15004 in the different experimental conditions.

It is also important to consider that the MTT assay measures cellular tetrazolium- reducing activity rather than directly determining cell number or neuronal survival [24]. Consequently, the changes observed in our experiments should be interpreted as alterations in MTT-reducing activity, which may reflect changes in cellular metabolic activity as well as cell viability.

A major finding of the present study is that pretreatment with ITH15004 improved MTT- reducing activity in NSC-34 cells exposed to O/R, ATP and nigericin. ITH15004 alone, at the concentrations tested, did not significantly modify MTT-reducing activity, suggesting that its effects under the experimental conditions were not associated with intrinsic cytotoxicity.

The improvement observed following ATP exposure is compatible with a potential contribution of purinergic receptor modulation. ITH15004 is a purine-derived compound previously described as a P2X7 receptor antagonist [21]. Since extracellular ATP can activate P2X7 receptors and promote cellular stress, antagonism of this receptor represents a plausible explanation for at least part of the protective effect observed in ATP-treated cells [17,18,21].

ITH15004 also improved MTT-reducing activity following O/R exposure. This observation is of particular interest because O/R directly interferes with mitochondrial respiratory function through inhibition of complexes I and V [19,20]. The improvement observed under these conditions suggests that the effects of ITH15004 may not be restricted to the modulation of extracellular ATP-dependent signalling. Nevertheless, the present experiments do not establish whether this response results from direct mitochondrial protection or from indirect effects on other cellular pathways.

The most pronounced effect was observed in nigericin-treated cells, in which ITH15004 restored MTT-reducing activity towards control levels. This finding suggests that ITH15004 can attenuate cellular damage induced by a stimulus that disrupts ionic homeostasis through a mechanism distinct from direct purinergic receptor activation. Taken together, the effects observed across the three experimental paradigms suggest that ITH15004 may influence cellular responses to different forms of stress. However, further studies are required to determine whether these effects share a common molecular mechanism.

Mitochondrial dysfunction and impaired energy metabolism are important features of ALS pathophysiology [7–9]. Motor neurons are particularly susceptible to disturbances in cellular energy metabolism, and mitochondrial abnormalities have been associated with oxidative stress, altered calcium regulation and activation of cell death pathways [8,9,15].

In our study, nigericin produced a marked reduction in intracellular ATP levels, whereas O/R and ATP did not produce a comparable significant reduction under the conditions tested. Importantly, ITH15004 pretreatment partially restored ATP levels in nigericin- exposed cells, increasing the measured signal from approximately 36.8% to 65.9% of control values.

This finding suggests that ITH15004 attenuates the reduction in cellular ATP associated with nigericin exposure. Nigericin alters potassium and proton gradients across biological membranes, potentially disturbing cellular and mitochondrial ion homeostasis [28]. Such ionic disturbances may contribute to impaired cellular energy balance. Nevertheless, the CellTiter-Glo assay measures cellular ATP content and does not directly determine mitochondrial ATP synthesis or respiratory activity. Therefore, the observed recovery should be interpreted as an improvement in intracellular ATP availability rather than direct evidence of restored mitochondrial oxidative phosphorylation.

The ability of ITH15004 to partially preserve ATP levels is consistent with its previously described pharmacological properties. In particular, previous work has shown that ITH15004 can modulate cellular processes involving mitochondrial calcium handling in chromaffin cells [22]. Although this previous observation provides relevant background for investigating the compound’s cellular actions, the present study does not establish a direct mitochondrial target for ITH15004. Additional experiments measuring oxygen consumption, respiratory capacity and mitochondrial membrane potential would be necessary to determine whether the improvement in ATP levels is associated with preserved mitochondrial bioenergetic function.

Mitochondria play a central role in the regulation of intrinsic apoptotic signalling. Under conditions of cellular stress, mitochondrial alterations can promote the activation of downstream caspases, including caspase-3, which participates in the execution phase of apoptosis [10,11]. Mitochondria-dependent apoptotic mechanisms have also been implicated in ALS, with previous studies reporting alterations in Bax, cytochrome c and caspase signalling in experimental models of the disease [26,27].

In our study, nigericin produced a pronounced increase in caspase-3/7 activity in NSC- 34 cells. This response indicates the activation of executioner caspases following nigericin exposure and is consistent with the involvement of apoptotic signalling in the cellular response to this stimulus. Importantly, ITH15004 pretreatment significantly reduced caspase-3/7 activity compared with nigericin treatment alone.

These results suggest that ITH15004 attenuates the activation of caspase-dependent cell death pathways under nigericin-induced stress. The reduction in caspase activity, together with the improvement in MTT-reducing activity and intracellular ATP levels, supports a protective effect of ITH15004 against nigericin-induced cellular damage.

Previous studies have demonstrated that mitochondrial dysfunction and cytochrome c- dependent apoptotic signalling contribute to neuronal injury in ALS models [10,26,27]. Although our results are compatible with the involvement of apoptotic pathways, the present experiments do not establish the precise sequence of molecular events leading to caspase activation or demonstrate that ITH15004 directly inhibits a specific component of the apoptotic machinery.

Calcium homeostasis is essential for neuronal function, and disturbances in intracellular calcium regulation have been associated with motor neuron vulnerability in ALS [9,15]. Sustained changes in cytosolic calcium can influence multiple cellular processes and may contribute to neuronal dysfunction under conditions of oxidative and mitochondrial stress.

Using the ratiometric fluorescent indicator Fura-2, we observed that nigericin induced an increase in cytosolic calcium concentration in NSC-34 cells. This response was concentration-dependent, with the largest increase observed at 15 µM nigericin. These findings demonstrate that nigericin exposure is associated with substantial changes in cytosolic calcium levels in this cellular model.

In the sequential perfusion experiments, ITH15004 alone produced a relatively small increase in cytosolic calcium. Subsequent addition of nigericin in the presence of ITH15004 produced a further increase, followed by a more pronounced response after ATP application. The ATP-induced response was included as a positive control to confirm that the cells remained responsive and that the experimental preparation and calcium-recording system were functioning appropriately.

These experiments provide evidence that ITH15004 and nigericin are associated with changes in cytosolic calcium dynamics. However, the experimental design does not allow us to conclude that ITH15004 prevents or reverses the calcium response induced by nigericin. Furthermore, Fura-2 measurements reflect cytosolic calcium changes and do not distinguish between calcium entry across the plasma membrane and calcium mobilization from intracellular stores.

The relevance of these observations lies in the established association between altered calcium homeostasis and motor neuron vulnerability in ALS [9,15]. Nevertheless, further experiments will be needed to clarify the mechanisms underlying the calcium responses observed in the presence of ITH15004 and nigericin.

Mitochondrial proteins involved in the regulation of energy metabolism and apoptotic signalling are altered in several models of neurodegeneration, including ALS [7–9,26,27]. Cytochrome c is an essential component of the mitochondrial electron transport chain and also participates in intrinsic apoptotic signalling following its redistribution from mitochondria to the cytosol [10,11,26]. VDAC1 is an outer mitochondrial membrane protein commonly used as a marker of mitochondrial content.

In the present study, Western blot analysis showed that nigericin reduced cytochrome c protein abundance relative to GAPDH. A similar pattern was observed for VDAC1. In contrast, ITH15004 treatment alone increased the abundance of both proteins, while co-treatment with ITH15004 and nigericin counteracted the reductions observed following nigericin exposure. In addition, the cytochrome c/VDAC1 ratio was reduced by nigericin and increased following ITH15004 treatment, with co-treatment restoring this ratio towards control values.

These observations suggest that ITH15004 modifies the abundance of mitochondrial- associated proteins and counteracts the changes induced by nigericin. The increase in cytochrome c and VDAC1 following ITH15004 exposure may be consistent with changes in mitochondrial protein content or mitochondrial abundance. However, the present data do not demonstrate mitochondrial biogenesis, which would require additional evidence, such as measurements of mitochondrial DNA content, other mitochondrial markers and functional assessment of mitochondrial mass.

An important methodological consideration is that the Western blot experiments were performed using total cellular lysates rather than separately isolated cytosolic and mitochondrial fractions. Therefore, the reduction in total cytochrome c abundance following nigericin exposure cannot, by itself, be interpreted as evidence of cytochrome c release into the cytosol or mitochondrial outer membrane permeabilization. Similarly, the restoration of cytochrome c abundance by ITH15004 does not directly demonstrate inhibition of cytochrome c release. Nevertheless, the changes in cytochrome c and VDAC1, together with the effects on ATP levels and caspase-3/7 activity, suggest that ITH15004 influences cellular responses associated with mitochondrial stress.

P2X7 receptors have been implicated in neuroinflammatory signalling and neuronal injury, making them potential therapeutic targets in ALS [17,18]. Activation of these receptors by extracellular ATP can induce ionic disturbances and promote downstream inflammatory and cell death pathways [12,18]. In this context, the expression of P2X7 receptors detected in NSC-34 cells and the reported antagonist activity of ITH15004 provide a rationale for investigating the contribution of purinergic signalling to the effects observed in our model [21].

The improvement in MTT-reducing activity following ATP exposure is compatible with a possible contribution of P2X7 receptor antagonism. However, the present study did not directly assess receptor function or demonstrate that P2X7 blockade is responsible for the protective effects of ITH15004.

Moreover, the protective effect observed against nigericin-induced damage suggests that additional mechanisms may contribute. Nigericin alters ion gradients through its ionophore activity and can induce potassium efflux independently of direct P2X7 receptor activation [25,28]. Consequently, the effects of ITH15004 against nigericin cannot be attributed solely to P2X7 receptor antagonism.

One possibility is that ITH15004 influences cellular responses downstream of, or independent from, purinergic receptor activation. Its previously described effects on cellular processes involving mitochondrial calcium handling [22] provide a rationale for further investigation, although the present study does not establish this as the mechanism underlying its protective effects. Future experiments using selective P2X7 receptor antagonists, receptor knockdown approaches and direct assessment of mitochondrial function will be important Further studies should therefore investigate mitochondrial respiration, membrane potential, reactive oxygen species production and mitochondrial content using complementary approaches. In addition, experiments specifically addressing P2X7 receptor function will be necessary to establish the contribution of purinergic signalling to the pharmacological effects of ITH15004. Validation in more complex cellular systems and relevant ALS animal models will also be required to determine the potential translational relevance of these findings.

In conclusion, the present study provides evidence that ITH15004 attenuates cellular damage induced by different stressors in NSC-34 motor neuron-like cells. Its effects are particularly evident following nigericin exposure, where ITH15004 improves MTT- reducing activity, partially restores intracellular ATP levels and reduces caspase-3/7 activation. In addition, ITH15004 modifies cytosolic calcium responses and counteracts nigericin-associated changes in cytochrome c and VDAC1 protein abundance. Although the precise mechanisms remain to be established, these findings support further investigation of ITH15004 as a potential modulator of cellular responses to mitochondrial and ionic stress relevant to ALS.

## SUPLEMENMTARY DATA

**S1.**
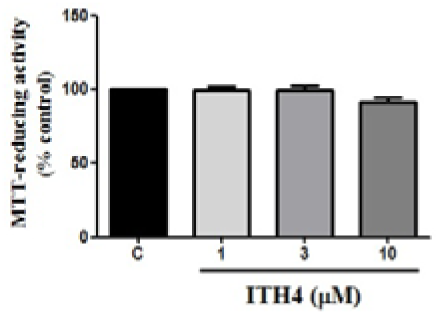
Effect of ITH15004 on cell viability NSC-34 cells were treated for 24 hours, with increasing concentrations of ITH15004 (1, 3, and 10 µM). Cell viability was determined by the MTT assay. Data are expressed as mean ± SEM from four independent cultures (N = 4) with three replicates per condition (n = 12). The treatment is displayed at the top. *One-way repeated-measures ANOVA, followed by* Tukey multiple comparison test post-hoc; no significant differences were detected (p > 0.05).

**S2.**
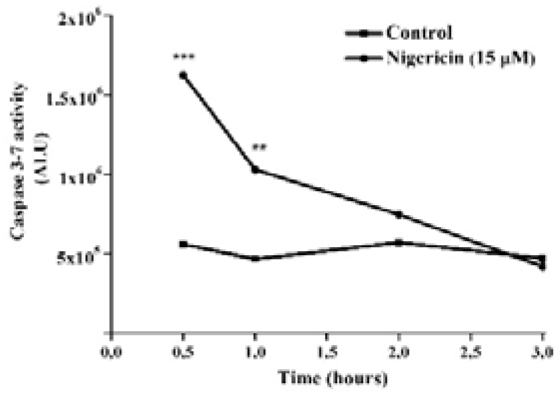
***Luminescence assay to establish the optimal incubation time for caspase-3/7 activity in NSC-34 cells.*** Cells were seeded into 96-well plates (50,000 cells/well) and treated for 24 h with Nigericin (15 µM). Luminescence was measured at different time points (0.5–3 h) using the Caspase-Glo® 3/7 assay kit (Promega). Data represent mean ± SEM of three independent experiments performed in triplicate (n = 9). One-way ANOVA analysis; p < 0.05 was considered statistically significant.

